# Low-Intensity Ultrasound Lysis of Amyloid Microclots in a Lab-on-Chip Model

**DOI:** 10.1101/2025.03.10.642458

**Authors:** Reza Rasouli, Brad Hartl, Soren Konecky

**Affiliations:** Openwater, San Francisco, California, United States

## Abstract

Amyloid fibrin(ogen) microclots are misfolded protein aggregates with β-sheet structures that have been associated with Long COVID and numerous thrombo-inflammatory diseases. These microclots persist in circulation and obstruct microvasculature, impair oxygen transport and promote chronic inflammation. Conventional thrombolytic therapies such as recombinant tissue plasminogen activator (rtPA) show limited efficacy against these aggregates due to their structure and composition. In this study, we assess the impact of low intensity focused ultrasound (LIFU) stimulation on amyloid microclot fragmentation, the role of cavitation in this process and investigate whether microbubble-assisted ultrasound can enhance their lysis. Amyloid microclot models were generated using freeze-thaw cycles followed by incubation. Microclots were exposed to ultrasound waves at 150 kHz, 300 kHz, 500 kHz, and 1 MHz under four conditions: ultrasound alone (US), ultrasound with microbubbles (MB + US), ultrasound with rtPA (rtPA + US), and ultrasound with both microbubbles and rtPA (MB + rtPA + US). Low-frequency ultrasound at 150 kHz resulted in a significant clot lysis with up to three-fold reduction in both clot size and the number of large clots. The addition of microbubbles enhanced clot lysis at 150 kHz, 300 kHz, and 500 kHz. These findings suggest that ultrasound, particularly at 150 kHz, is a promising method for amyloid microclot lysis. The combination of ultrasound with microbubbles and rtPA further improved clot fragmentation, rendering it a potential therapeutic tool for conditions like Long COVID.

## Introduction

Protein misfolding into abnormal amyloid aggregates is a defining hallmark of a wide array of human disorders, known as amyloidoses.^1^ Over 50 diseases, including neurodegenerative conditions (e.g., Alzheimer’s, Parkinson’s, Huntington’s diseases), metabolic/endocrine diseases (e.g., Type II diabetes), and systemic amyloidoses (e.g., light-chain amyloidosis), are characterized by pathological amyloid accumulation.^1,2^

In these disorders, soluble proteins misfold and aggregate into insoluble fibrillar aggregates where they lead to disruption of tissue architecture and function. These protein aggregates not only can interfere with cellular processes but also lead into disease progression through inducing chronic inflammation and oxidative stress. ^1,2^ Although the composition of amyloid microclots varies in different biological conditions, their common structural feature is the formation of ordered β-across-β architecture. ^3,4^ The cross-β core increases the stability of amyloid fibrils, rendering them insoluble and proteolysis-resistant.^2^

Among amyloid-related pathologies, amyloid fibrin(ogen) microclots have recently gained attention due to their role in Long COVID (post-acute COVID-19 syndrome) and cardiovascular diseases. Recent studies have identified amyloid fibrin(ogen) microclots in the blood of patients with Long COVID. ^3,5–8^ These microclots have been detected as a central pathological factor which are associated with persistent symptoms such as fatigue, breathlessness, and cognitive dysfunction. Coagulation of such aggregates in the microvasculature can block capillary flow and impair oxygen transport to tissues, which in turn lead to hypoxia, chronic organ dysfunction and inflammation.^8,9^ Proteomic analyses showed that COVID-related microclots contain pro-inflammatory molecules and clotting factors that result in persistent inflammation and resistance to fibrinolysis.^7^

Current therapies for Long COVID focus primarily on enzymatic clot dissolution, where fibrinolytic enzymes such as rtPA act as a serine protease to convert plasminogen to plasmin and degrade fibrin networks. Although enzymatic therapies are effective against conventional thrombi, they struggle to address amyloid-rich microclots due to their structural resistance to enzymatic breakdown. This fibrinolysis resistance is further exacerbated in COVID-19 patients by upregulation of α-(2)-antiplasmin which inhibits plasmin activity and limits the effectiveness of rtPA. ^6,10^ Clinical observations confirm this inefficiency, where COVID-19 patients often showed poor response to tPA-mediated thrombolysis.^10^

Another strategy to treat circulating microclots is filtration-based strategies to physically remove them from the bloodstream. Apheresis is reported for severe COVID patients,^11^ in which blood is drawn from the patient, passes through an extracorporeal filter, and then returns to the body. However, extracorporeal methods are invasive, resource-intensive, and involve health risks such as bleeding and infection which limit their application to acute conditions or experimental settings. Given the pathological impact and therapy resistance of amyloid microclots, there is a clear need for innovative and non-invasive treatments that can effectively disrupt circulatory amyloid microclots in patients and restore microvascular function.

Ultrasound has a long successful history in treating blood clots in clinical settings. Addition of transcranial ultrasound exposure to IV tPA treatment significantly improved thrombolysis in a phase II trial for acute ischemic stroke, leading to early arterial recanalization and enhanced clinical recovery rates compared to tPA alone.^12^ In deep vein thrombosis (DVT), ultrasound-assisted catheter thrombolysis outperformed enzymatic thrombolysis alone with high clot clearance (around 70% complete lysis, 91% partial/complete lysis) and faster treatment.^13^

There is a strong theoretical and clinical basis for employing ultrasound to disrupt microclots.^14–18^ Ultrasound in the low-kilohertz to low-megahertz range can induce mechanical phenomena such as acoustic streaming, cavitation, shear forces, and radiation forces that can be used to promote clot fragmentation.^19^ Acoustic streaming generates convective currents and shear stress around the thrombus, enhancing penetration and mixing of fibrinolytic agents into the clot.^14,20^ Acoustic streaming and acoustic radiation force (ARF) can also physically erode clot structure by pushing fluid through the fibrin mesh, loosening pore spaces and increasing permeation of plasminogen activators into the thrombus.^18^ This streaming action also exposes new fibrin binding sites to plasmin, enhancing enzymatic breakdown.^18^

Numerous *in vitro*, animal, and clinical studies report faster and more effective thrombolysis when ultrasound is applied alongside fibrinolytic drugs.^18^ Unlike chemical thrombolysis which relies solely on biochemical mechanisms, ultrasound offers advantage by offering a mechanical mode of action that can help to overcome the fibrinolysis resistance of amyloid clots.^21^ The combination of acoustic streaming and cavitation can physically disrupt and weaken the clot. Prior studies on clots have shown that ultrasound can significantly accelerate clot lysis by mechanical forces.^17^ For example, exposure to 1 MHz ultrasound at ∼2 W/cm^2^ produced streaming with a shear rate on the order of 40 s^−1^, roughly doubling the enzymatic lysis rate *in vitro*.^16,22^ In microbubble-assisted sonothrombolysis, oscillating microbubbles create vigorous microstreaming and localized shear forces that significantly accelerate clot breakdown.^20,22^ Experimental studies have visually confirmed these effects, showing that stable cavitation bubbles preferentially erode clots and correlate with faster lysis rates compared to enzymatic treatments.^19,20^ Furthermore, microbubble-assisted sonothrombolysis facilitates cavitation to enhance clot lysis and significantly boost clot dissolution.^15^

Despite significant progress in sonothrombolysis for large vascular clots, no reported studies have specifically explored ultrasound-mediated lysis of circulating amyloid microclots which are recognized as a key therapeutic target in Long COVID. Here, we hypothesize that applying ultrasound can effectively disrupt amyloid fibrin microclots *in vitro* via acoustically induced streaming. To test this, we employed a LOC model that enables precise control over microclot exposure conditions, allowing systematic evaluation of the effects of frequency, shear stress, and microbubble presence on clot disruption. Our goal is to identify optimal ultrasound parameters for microclot fragmentation and lay the groundwork for noninvasive strategies to mitigate microvascular dysfunction and coagulopathies in amyloid-related pathologies such as Long COVID.

## Materials and Methods

### Materials

Porcine plasma was obtained from Innovative Research, Inc. (United States) and stored at -20°C until use. Plasma aliquots (1 mL) were used for microclot formation. rtPA was purchased from Sigma-Aldrich (MilliporeSigma, United States) and reconstituted according to the manufacturer’s protocol to achieve a final working concentration of 3 *μ*g/mL. Thioflavin T (ThT) was purchased from Sigma-Aldrich (St. Louis, MO, USA). Vevo MicroMarker® contrast agents (FUJIFILM VisualSonics Corp., Japan) were used as exogenous microbubbles and were diluted in PBS buffer per manufacturer protocol.

### Amyloid Microclot Formation

Amyloid fibrin(ogen) microclots were generated by subjecting porcine plasma to repeated freeze-thaw cycles, followed by incubation at 37 °C. Plasma aliquots (1 mL) underwent 12 freeze-thaw cycles, with each cycle consisting of freezing at -20°C and thawing at 37°C for two hours. The aliquots were diluted 1.5 x before the ultrasound treatment.

### Lab on Chip Device Fabrication

The LOC device was designed to replicate a 6 mm diameter popliteal vein, providing a controlled environment for targeted ultrasound exposure of amyloid microclots. The popliteal vein was selected due to its accessibility for transcutaneous ultrasound exposure rendering it a relevant site for potential clinical applications of ultrasound. The device molds were designed using Onshape (PTC Inc., United States) CAD software and fabricated via digital light processing (DLP) technology using a 285 D Printer (CADWorks3D, Canada). After UV treatment, Momentive RTV615 silicone elastomer (RS Hughes, United States) was cast into the molds and cured following manufacturer-recommended conditions to form the microfluidic channels. The cured elastomer was then bonded onto a 20 *μ*m silicone sheet (Gteek, United Kingdom) using plasma surface treatment.

### Experimental Setup

The schematic representation of the LOC platform, transducer arrangement, and acoustic bath setup is shown in Figure 1. The experimental setup consisted of a custom-designed and 3D printed acoustic water bath, which housed the LOC device, a piezoelectric transducer, and supporting components. The transducers were mounted at the bottom of the bath. The LOC chip was mounted on top of the water bath using a custom-designed holder, securing its position at the pre-measured focal plane of the ultrasound transducers to maintain consistent acoustic exposure across all experimental conditions.

**Figure 1.**
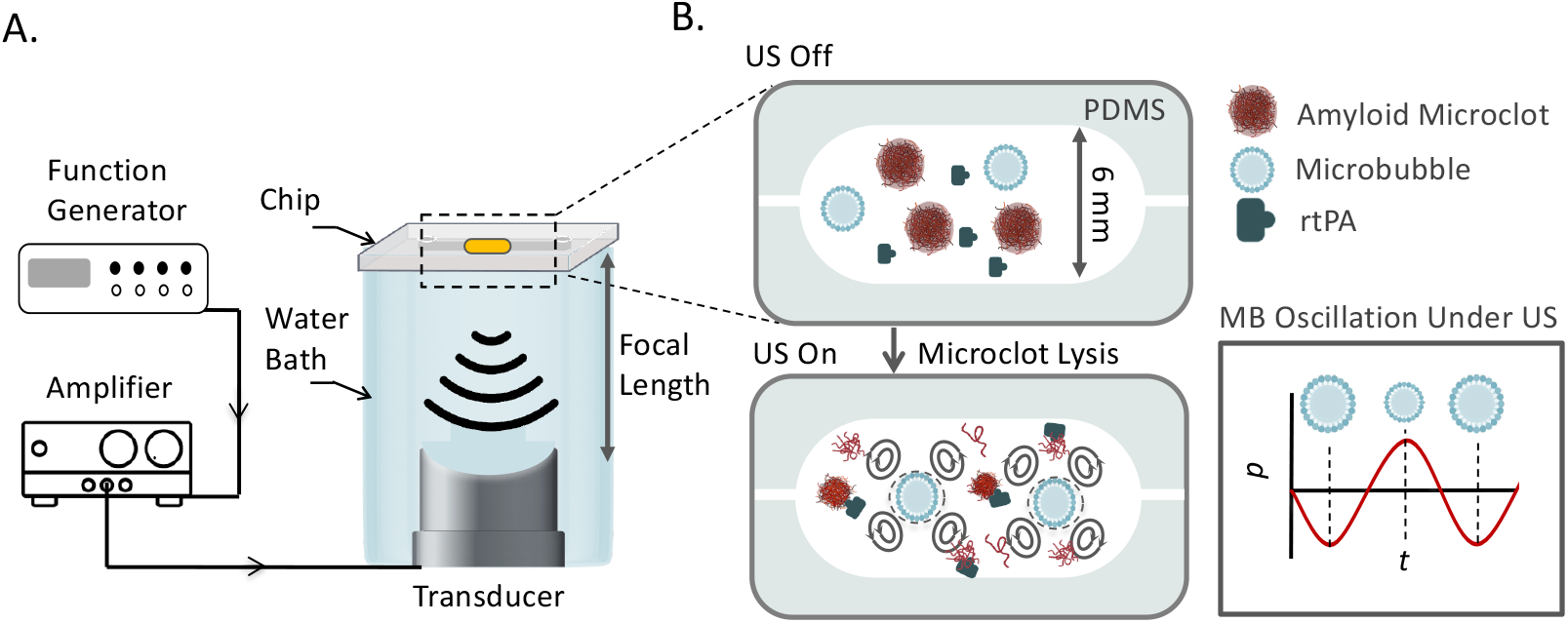
**A**. Schematic of the experimental platform for ultrasound-mediated amyloid microclot lysis. The LOC platform is placed in an acoustic water bath and exposed to focused ultrasound at 150 kHz, 300 kHz, 500 kHz, and 1 MHz. A function generator controls the ultrasound frequency, and an amplifier amplifies the signal. **B**. Zoomed-in view of the LOC channel, showing the interaction of ultrasound, microbubbles, and rtPA with microclots. Microbubbles undergo volumetric oscillations in response to acoustic pressure waves, generating streaming effect and shear stress to lyse microclots.

A function generator (DG4162, Rigol Technologies Co., Ltd., China) controlled the ultrasound frequency and waveform, while an amplifier (Model 1040L, Electronics & Innovation, Ltd., United States) regulated the voltage to the transducers. The bath was filled with degassed water as a coupling medium for ultrasound wave transmission while minimizing acoustic attenuation.

Experiments were conducted under static flow conditions in the LOC platform. Ultrasound was applied at four different frequencies: 150 kHz, 300 kHz, 500 kHz, 1 MHz. The mechanical index (MI) is defined as 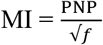, where PNP is the measured peak negative pressure (MPa), and *f* is the ultrasound frequency (MHz). The MI was maintained at 0.3 for all the experiments by adjusting the ultrasound pressure accordingly for all frequencies. The thin silicone membrane at the base of the chip allows acoustic waves to enter the microfluidic chamber while minimizing attenuation. Inside the LOC channel, ultrasound waves interact with the plasma and amyloid microclots, generating acoustic streaming, cavitation, and shear forces. These interactions induce microstreaming and localized mechanical forces, which contribute to the mechanical fragmentation and potential enzymatic enhancement of clot lysis.

### Microscopy and Image Analysis

Thioflavin T was used as a fluorescent dye to stain amyloid fibrils. A stock solution of 0.03 mM Thioflavin T was prepared in phosphate-buffered saline (PBS), pH 7.4. The solution was filtered using a 0.2 *μ*m syringe filter to ensure clarity. Samples were incubated with Thioflavin T at room temperature for 30 minutes prior to imaging.

Amyloid microclot lysis was evaluated based on clot count reduction and size reduction. The microclots morphology and count were monitored before and after ultrasound exposure, and changes in clot size were analyzed using fluorescence microscopy. Image analysis was performed using segmentation and particle detection plugins in ImageJ software, which measured clot area, diameter, and count. One-way ANOVA was used to compare clot lysis across experimental groups, with a significance threshold of p < 0.05.

## Results & discussion

### Amyloid Microclot Model Formation

Development of a reproducible amyloid fibrin microclot model is essential for investigating their role in clinical disorders including Long COVID, Alzheimer’s, type 2 diabetes.^3,4^ Various *in vitro* methods have been reported to produce amyloid microclots including drying, heating, exposure to acidic pH, hydrophobic surfaces, and organic solvents.^4^

In this study, we used the freeze-thaw method and incubation at biological temperature to induce amyloidogenesis. Freeze-thawing results in ice-water interfaces in plasma and introduces structural stress into proteins which could induce protein misfolding and aggregation.^23^ Moreover, the incubation at room temperature can promote amyloidogenesis through oxidative stress and spontaneous fibrinogen misfolding that ultimately enhances the formation of amyloid fibrin microclots. Freeze-thaw-based amyloid formation is a cell and enzyme-free approach. In this method, protein structural modification results from intrinsic protein dynamics rather than external biochemical interactions, offering reproducibility and control over aggregation conditions.^23,24^

To verify amyloid formation by freeze-thawing, we employed the Thioflavin T (ThT) dye, which selectively binds to β-sheet-rich amyloid fibrils and enhances fluorescence upon interaction. Fluorescent ThT staining is widely used in amyloid research as a reliable marker for detecting the amyloid transition of fibrinogen, owing to its high specificity for cross-β structures.^3,5,6,25^

Figure 2 A shows the fluorescence microscopy images of freeze-thawed plasma samples stained with ThT. In the earlier cycles, there are no visible ThT-labeled amyloid structures. However, green, fluorescent ThT-labeled particles emerge starting from cycles 7, indicating a gradual amyloid microclot formation. By cycle 8, a marked increase in ThT-tagged particles can be observed which shows significant amyloid microclot formation and with each cycle more amyloid microclots appear in the sample. Figure 2 B shows the quantitative clot counts as a function of cycles, confirming a significant increase in amyloid microclots after cycle 7. This cycle-dependent increase in amyloid microclots counts confirms amyloidogenesis by repeated freeze-thaw cycles and incubation. This experiment was repeated twice to ensure reproducibility. However, since plasma contains natural proteins, batch-to-batch variations are inherent. As a result, the data presented here is primarily aimed to show the trend of increasing clot formation with successive cycles rather than serving as an exact reference for clot numbers.

**Figure 2.**
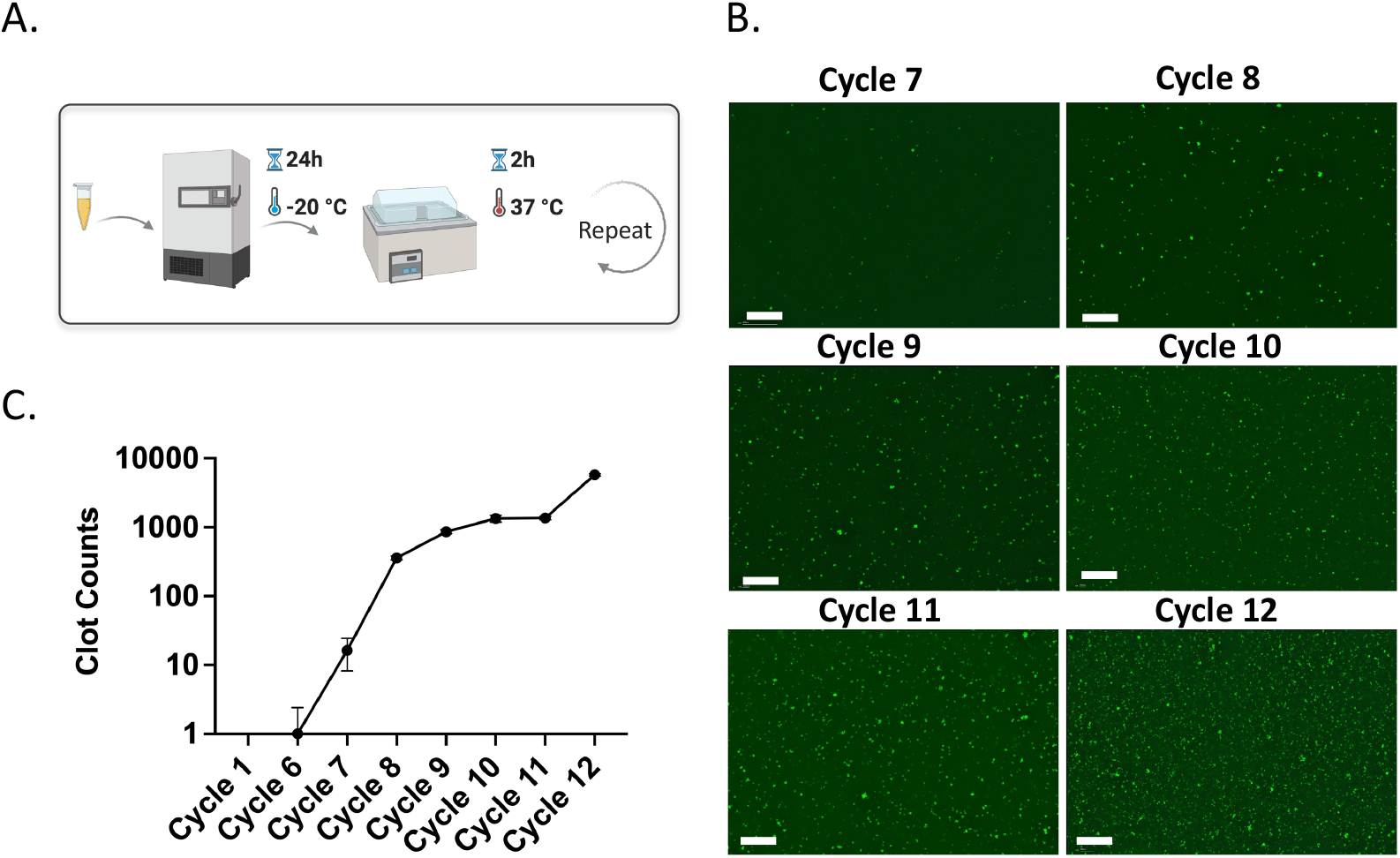
**A**. Schematic representation of amyloid microclot model formation using freeze-thaw cycles. Fluorescence microscopy images of ThT-stained porcine plasma across freeze-thaw cycles. Scale bar is 200 µm. **C**. Clot count quantification graph showing the increase in amyloid microclots over cycles.

### Effect of Ultrasound on Amyloid Microclots Across Different Frequencies

After amyloid microclot formation, we explore how different ultrasound frequencies impact both microclot morphology and count. The different mechanical interactions of ultrasound with microclots are frequency-dependent, as acoustic forces such as acoustic radiation force and acoustic streaming exhibit varying levels of dominance based on their inherent frequency ranges.^14^ The balance between these forces dictates whether ultrasound leads to microclot fragmentation, which is our aim, or instead results in displacement or even aggregation of the microclots. Hence, identifying the optimal frequency and experimental conditions for effective clot disruption is crucial for the potential therapeutic applications of our ultrasound platform. To this end, we tested the effect of different ultrasound frequencies on the lysis of microclots by subjecting them to 150 kHz, 300 kHz, 500 kHz, and 1 MHz.

This experiment was also repeated in the presence of rtPA, a serine protease used as a thrombolytic agent to dissolve blood clots. rtPA works by binding to fibrin in microclots and then converting plasminogen into plasmin, which subsequently dissolves the clot through a process known as fibrinolysis. We sought to determine whether ultrasound increases the rate of clot dissolution when rtPA is used and whether a possible synergistic effect of the acoustic forces and pharmacological thrombolysis can be observed.

Figures 3A and 3B show fluorescence microscopy images of ThT-tagged microclots treated with different ultrasound (US) frequencies for US-only and US+rtPA groups. The control samples show a high density of large (over 30 microns) amyloid aggregates, which are uniformly distributed across the image. Upon exposure to 150 kHz waves, the microclots exhibit a marked reduction in size. At higher frequencies, however, less fragmentation can be observed, and the clots remain intact.

**Figure 3.**
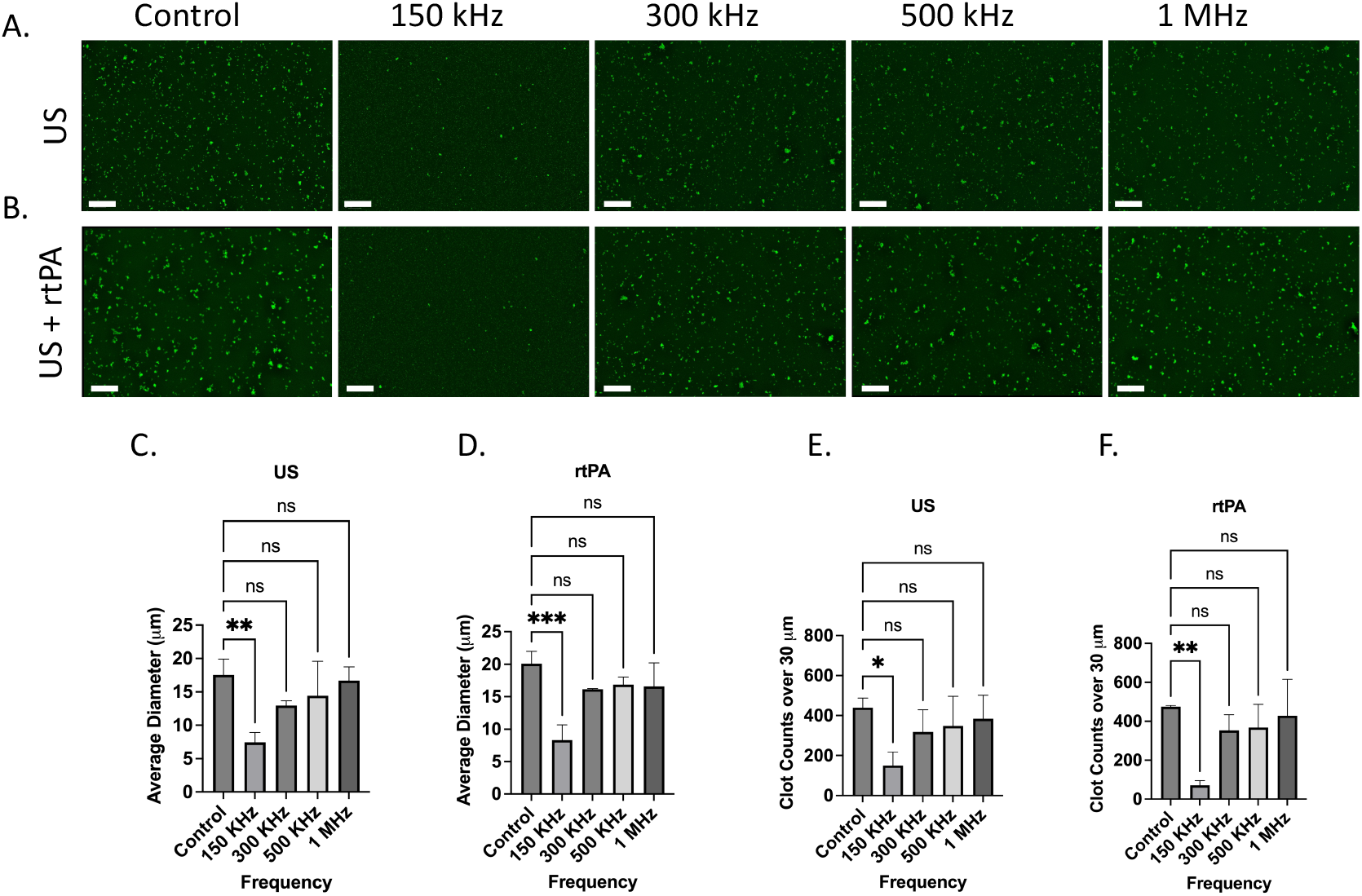
Effect of ultrasound (US) and ultrasound combined with rtPA (US + rtPA) on microclot lysis at different frequencies. **A, B**. Fluorescence microscopy images of microclots under **A**. ultrasound alone and **B**. ultrasound with rtPA at control, 150 kHz, 300 kHz, 500 kHz, and 1 MHz. C, D. Bar graphs showing average clot diameter as a function of ultrasound frequency for **C**. US-only and **D**. US + rtPA conditions. Clot size is significantly reduced at 150 kHz, with a minor reduction at higher frequencies. **E, F**. Clot counts for aggregates larger than 30 µm across different conditions, demonstrating a frequency-dependent clot fragmentation. Error bars represent standard deviations. Scale bar is 200 µm.

To quantify microclot lysis under US treatment, image analysis was performed, and the average clot diameter and count of clots above 30 *μ*m at each testing condition were calculated for three samples to ensure statistical robustness. Figure 3C, D show the average clot diameter as a function of ultrasound frequency for both US and US+rtPA samples. In agreement with the microscopy images, a significant reduction in clot size is observed at 150 kHz, while exposure to higher frequencies leads to less pronounced fragmentation. US+rtPA shows a similar trend of reduction in average diameter and large clot count at 150 kHz. For US+rTPA the mean clot diameter decreases from ∼20 *μ*m in the control to ∼8 *μ*m at 150 kHz, confirming significant fragmentation at this frequency. At 300 kHz and higher frequencies there is no statistically significant change in clot size for both US and US+rTPA.

The findings indicate that 150 kHz ultrasound is the most effective frequency for reducing both large clot count and size in both US-only and US+rtPA groups, whereas higher frequencies (300 kHz-1 MHz) result in less pronounced disruption. One potential explanation for clot lysis at 150 kHz is the formation of endogenous microbubbles, which can result in boundary-layer-driven acoustic microstreams.^14^ During the 150 kHz experiment, bubbles became visibly present, whereas no such bubble formation was observed at higher frequencies. Plasma and most biological fluids naturally contain gas nuclei that act as precursors for bubble formation under ultrasound exposure. Upon formation, these bubbles oscillate in response to ultrasound actuation and create localized streaming which in turn can enhance clot lysis.

### The Role of Cavitation in Ultrasound-Induced Clot Fragmentation

To investigate whether the mechanism of fragmentation at 150 kHz was directly linked to cavitation and acoustic streaming, we compared the effect of ultrasound exposure on degassed and non-degassed plasma samples. Cavitation requires dissolved gas nuclei to form microbubbles and thus degassing the plasma minimizes endogenous cavitation and bubble formation.

Figure 4 shows the fluorescence microscopy results of microclots in degassed and non-degassed plasma. Clot lysis and reduced micro clot size was observed in the gassed plasma sample after treatment with 150 kHz acoustic waves as shown by the images in Figure 4A. The same ultrasound waves failed to lyse microclots in degassed plasma samples and they remained largely intact. The absence of visible bubble formation in the degassed sample during the test as compared to the non-degassed sample suggests that clot lysis at 150 kHz can be attributed to the formation of endogenous bubbles and their attendant acoustic streaming.

**Figure 4.**
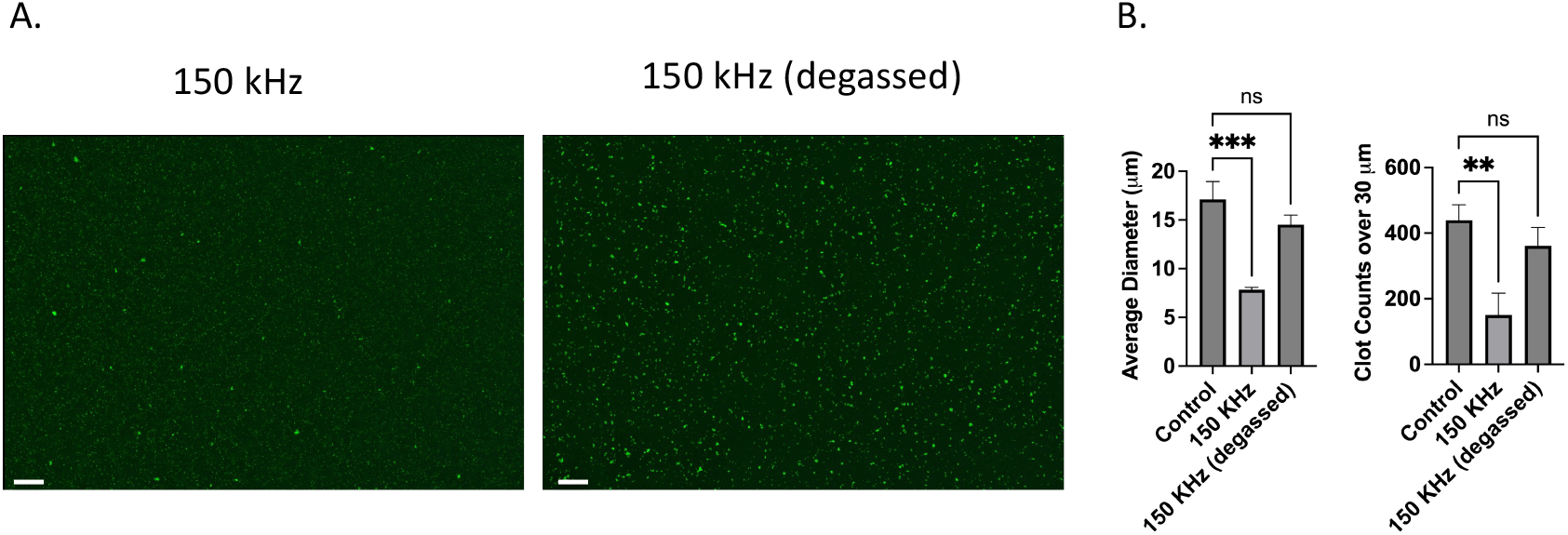
Effect of degassing on ultrasound US-induced microclot lysis at 150 kHz. **A**. Fluorescence microscopy images of microclots exposed to 150 kHz ultrasound in non-degassed (left) and degassed (right) conditions. Clot size remains larger in the degassed condition, indicating reduced lysis efficiency. Scale bars represent 100 *μ*m. **B**. Quantitative analysis of clot size and count. The left graph shows the average clot diameter across conditions.150 kHz stimulation of non-degassed sample significantly reduced clot size compared to the control, while the degassed condition retains larger clots. The right graph displays the number of clots over 30 *μ*m, showing a significant reduction for non-degassed, whereas degassed conditions result in higher clot retention.

Quantitative data on microclot size after ultrasound exposure is depicted in Figure 4B for both degassed and non-degassed samples. The left graph investigates microclot lysis in degassed vs non degassed plasma where cavitation effects are inhibited. The y-axis is the mean clot diameter, and the x-axis is frequency. Although the degassed samples showed a minor decrease in clot size it is considered statistically nonsignificant with diameters ranging from ∼17 to ∼14.5 *μ*m compared to ∼8 *μ*m in non-degassed sample. Moreover, the number of large clots over 30 *μ*m decreased for non-degassed sample to approximately a third of the control while in the degassed sample the variation was not statistically significant. This supports the hypothesis that clot lysis observed at 150 kHz is due to the cavitation, likely through cavitation-induced acoustic streams.

### 3.3 Effect of Microbubbles and rtPA on Clot Lysis Across Frequencies

Previous results showed that cavitation is crucial for clot fragmentation at 150 kHz. In this section, we introduce exogenous microbubbles (MBs) into the plasma to promote stable cavitation. We use gas-filled microbubbles ∼1-2 *μ*m in diameter which are coated with a phospholipid shell to render them more stable. In response to acoustic waves, these microbubbles undergo rapid compression and expansion cycles and generate localized fluid motion known as acoustic streams. Unlike endogenous cavitation, which relies on the presence of dissolved gas nuclei that varies between samples, contrast agents provide more consistent and controlled bubble dynamics and acoustic streaming. We hypothesize that the use of exogenous bubble can further increase lysis. In this section we also investigate the impact of MBs in combination with rtPA on clot lysis to assess whether this approach would further enhance enzymatic clot degradation.

Figure 5 shows the fluorescence microscopy images of ThT-stained plasma in the MB and MB + rtPA groups at the tested frequencies. In both groups, clot fragmentation is more pronounced at 150 kHz and 300 kHz, showing a significant reduction in clot size. Fragmentation is also observable with less impact at 500 kHz, while at 1 MHz, the effect diminishes significantly. Clot size analysis shown in Figure 5 further supports these observations and confirms a substantial decrease in clot size. The average clot diameter significantly decreases from ∼17 *μ*m in the control group to ∼7.5 *μ*m at 150 kHz, ∼8.5 *μ*m at 300 kHz, and ∼9 *μ*m at 500 kHz, while at 1 MHz the clot size remained about 10 *μ*m. Additionally, the large clot count decreased from ∼450 in the control to ∼70 at 150 kHz, ∼120 at 300 kHz, ∼180 at 500 kHz, and ∼350 at 1 MHz. The results confirm the capability of the ultrasound system in combination with MBs for fragmentation and elimination of large microclots.

**Figure 5.**
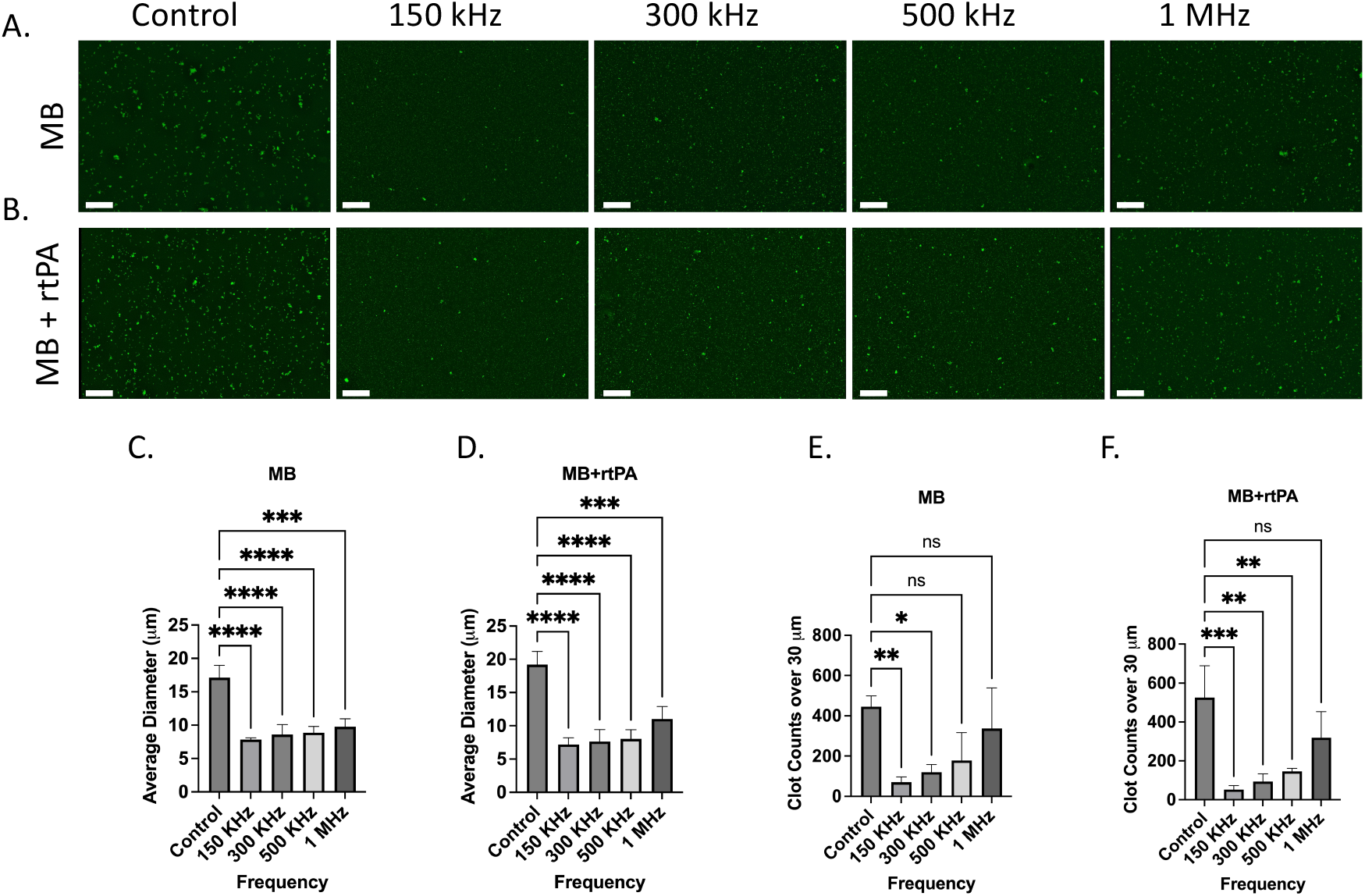
Effect of acoustically stimulated microbubbles (MB) and microbubbles combined with rtPA (MB + rtPA) on microclot lysis at different frequencies. **A, B**. Fluorescence microscopy images of microclots under **A**. ultrasound and Microbubbles and **B**. combination of ultrasound, microbubbles with rtPA at control, 150 kHz, 300 kHz, 500 kHz, and 1 MHz. C, D. Bar graphs showing average clot diameter as a function of ultrasound frequency for **C**. MB **D**. and MB + rtPA conditions. **E, F**. Clot counts for aggregates larger than 30 *μ*m. Scale bar is 200 *μ*m.

A similar trend, with a slightly stronger effect, can be seen in the MB + rtPA conditions, where the average clot diameter decreases from ∼ 19 *μ*m in the control to ∼ 7 *μ*m at 150 kHz, ∼ 7.5 *μ*m at 300 kHz, and ∼ 8 *μ*m for 500 kHz. The large clot count declines from ∼ 525 in the control group to ∼50 at 150 kHz, ∼ 90 at 300 kHz, ∼ 150 for 500 kHz, while the count remained over 300 for 1 MHz. The observed reduction in clot size and count suggests a potential synergistic interaction between ultrasound, microbubbles, and enzymatic lysis, necessitating further investigation to optimize this effect. This is particularly relevant given that acoustic streaming is reported to be one of the strongest mechanisms to induce normal advection and enhancing mixing.^26^ Strong and rapid mixing can accelerate enzymatic clot degradation by facilitating the binding of rtPA to fibrin within the clot, thereby promoting the conversion of plasminogen to plasmin.

These results indicate that microbubble-assisted ultrasound stimulation is more effective for clot fragmentation at lower frequencies (150–500 kHz). The addition of exogenous contrast agents enhances cavitation and acoustic streaming, resulting in more effective clot disruption. Also, in MB added groups the clot fragmentation was observed across multiple frequencies, while in the US-only group, lysis was observed only at 150 kHz. These results emphasize the potential of MB-assisted ultrasound therapy for targeting persistent amyloid microclots in conditions such as Long COVID.

## Conclusion

In this study, we showed that ultrasound stimulation is an effective strategy for lysing amyloid fibrin(ogen) microclots, particularly at 150 kHz. Using a LOC model, we observed a significant reduction in clot size and large clot count after ultrasound exposure. The lack of fragmentation in degassed plasma confirms that cavitation plays a key role in clot lysis and enhancing fibrinolysis. Microbubble contrast agents further improved clot fragmentation through stable cavitation and generating acoustic streams. The combination of ultrasound microbubbles and rtPA (MB + rtPA + US) resulted in the largest reduction in clot size. This suggests that mechanical disruption enhances enzymatic penetration and fibrinolysis. These findings support the potential of ultrasound-enhanced therapy for amyloid-associated coagulopathies where persistent microclots impair vascular function. Ultrasound-based treatments could serve as a non-invasive tool to restore blood flow and improve oxygenation in affected patients. Future research should focus on refining ultrasound parameters for *in vivo* use, evaluating safety and efficacy, and exploring its potential with thrombolytic drugs.

## Acknowledgment

The authors would like to thank Dr. Muhammad Zubair for his contributions to the measurement and characterization of the focal length of the ultrasound transducers.

